# A reduced-memory multi-layer perceptron with systematic network weights generated and trained through distribution hyper-parameters

**DOI:** 10.1101/2022.03.01.482493

**Authors:** Neha Vinayak, Shandar Ahmad

## Abstract

A multi-layer perceptron (MLP) consists of a number of forward-connected weights (*W*_*ijk*_) from each feeding layer node (*n*_*ij*_) to the many initially equivalent nodes (*n*_*i*+1,*k*_) in the next layer. Exact *a priori* order and search space of these weights (*W*_*ijk*_) is random and prone to redundancy, irreproducibility and non-optimality. We demonstrate that a weight subspace (*W*_*ijk*_ for each *i* and *j*), generated systematically using a statistical distribution with predefined breakpoints and Genetic algorithm-trained hyper-parameters substantially reduces the computational complexity of an MLP and produces comparable or better performance than similarly trained equivalent models with fully defined weights. This distribution based neural network (DBNN) provides a novel framework to create very large neural network models with currently prohibitive memory requirements.

## 1. Introduction

Multi-Layer Perceptrons (MLP) solve ubiquitously wide range of computational problems ranging from computer vision (Mas and Flores [14]), medical sciences (Amato et al. [1]), to business marketing (Tkáč and Verner [23]) and robotics (Nanda et al. [16], Besari et al. [3]).

One of the barriers in the seamless application of MLP to complex problems is the need for large memory, as the number of weights to be optimized increases rapidly with large hidden layers and feature-rich models. Number of efforts have been recently made to solve this problem by defining alternative approaches to initialize and train network paramers. For example, Courbariaux et al. [4] proposed training of weights using a binary −1,+1 or ternary −1, 0, +1 weight system. The training was performed by backpropagating the errors and training the binary weights instead. Shayer et al. [22] suggested another way to compute the gradient for the binary weights. They used the local reparametrization method by Kingma et al. [11] to compute the gradients assuming that the pre-activation values for layer *i* are approximately Gaussian.

Binary space neural network models are indeed highly simplified and look for a solution in a much reduced space quickly. However, the actual search space is limited and is intuitively likely to miss an optimal solution and therefore alternative methods of defining neural network weight assignments with low explicit dimensionality, which can search solutions of complexity comparable to the standard randomly initialized models will be very useful as an alternative MLP framework.

Any indirect assignment of MLP weights renders the parameter space non-differentiable and hence alternatives to backpropagation training are needed together with any new approaches to define connection weight space. It may be noted that an exact mechanism how the neural network parameters actually learn is a complex problem. Some researchers in the field have provided useful insights into the MLP training process. For example, Jesus et al. [9] have observed that the final set of weights lie in the locality of the initial randomly initialized ones. In fact, in cases where this observation was true, the model was found to be trainable and concluding at the nearest local minima. On the other hand, if the final weights were found to be spatially far from the initial set, they observed that the model remained untrained. These observations highlight the challenges in using backpropagation model on standard models of MLP and hence the need to find better methods of initializing weights and arriving at solutions far from the local hyperspace defined by random initialization.

In Goodfellow et al. [7] many method of initializing weights for a Multi Layered Perceptron (MLP) have been discussed. The most general and widely used methods are randomly allocating weights from a normal or uniform distribution. Other suggested methods are using normalized initialization Glorot and Bengio [6] of the complete weight matrix using a suggested formula for normalization based on the number of nodes in the source and destination layers. This helps to choose an initial scale for the weights.

Methods like random orthogonal initialization Saxe et al. [20] and sparse initialization Martens [13] have also been suggested. All these methods use the same scale for the entire weight matrix. Scale refers to the spread of the distribution, also denoted by its standard deviation (*σ*).

Among all these works, use of distribution-generated or systematically defined weights has been made mostly for the purpose of initialising the weights of an MLP.

- One architecture which uses distribution based weights is Bayesian Neural Networks by MacKay [12]. It replaces each single weight of the neural network with a distribution to allow for probabilistic measures but does not reduce the number of trainable parameters.
- Another architecture which works on the principle of using weights generated by another system is HyperNetworks by Ha et al. [8]. The weights of a HyperNetwork architecture are generated by another network, called HyperNetwork, which sits over the main network. The weights of this HyperNetwork are trained, leading to a reduced search space and less number of trainable parameters. The authors show that Hypernetworks perform well for RNN, LSTM and CNN. We have implemented HyperNetwork for training a standard MLP and compared its performance with our proposed model.

To explore MLP architectures with high performance and low cost, Kim et al. [10] have proposed Network Architecture Search (NAS). They have classified NAS into neuro-evolutionary algorithms (like Genetic Algorithm), reinforcement learning based algorithms (like CNN, RNN) and One-shot architecture search (like HyperNetworks, Parameter sharing). The complete process of network architecture search involves the following: a) defining the search space, b) search strategy and c) evaluation of results.

To the best of our knowledge, none of the available works investigate if the generation process of these weights could also be used as part of the training protocol, whereby the large space of neural network parameters is trained by way of finding the best hyper-parameters of a generative model such as a Gaussian distribution, which also produces a trained MLP. In this work, we explore the feasibility of such an approach to define the search space through a model named as Distribution based Neural Network (DBNN). It uses a per-node statistical distribution to initialise an MLP as well as train it by finding its best hyper-parameters. None of the above architectures exploit the fact that there is architectural equivalence of weights connecting a node to its forward layer. DBNN assigns weights emerging from each node by substituting the random values using a distribution with pre-defined breakpoints. The training is achieved by allowing these breakpoints to scan a search space that can generate the optimum network weights. As a search strategy for DBNN, we explored the backpropagation and genetic algorithm techniques.

In summary, we propose DBNN as a one-shot architecture search space definition technique with a neuro-evolutionary search strategy that represents values of all outgoing weights connected to a given node *n*_*ij*_ being generated by a statistical distribution *D*(*p*_1_, *p*_2_, ..*p*_*h*_). An entire MLP model is completely defined by a set of these weight matrices, each of which is generated by its corresponding initial or trained hyper-parameters. Thus, instead of the complete weight matrix, only the distribution hyper-parameters need to be trained. So, now there are *m* such distributions for layer *i* and these may not all be on the same scale. For an MLP with K connections on the average between a layer *i* and *i* + 1, the complexity of the model is reduced by *K/H*, where *H* is the number of hyper-parameters through which weights are defined and selected from a distribution.

The feasibility of this proposed DBNN model has been evaluated in the following terms:

- First, we assess if the model is trainable. Since, the weights are generated from a distribution and the distribution hyper-parameters are trained, backpropagation of errors cannot be applied for training. Experiments have been conducted with gradient descent based search strategy, but results observed were not good. So, we have explored other optimization methods to train our model. Genetic algorithm based optimization is one such method which is used to optimise the parameters (weights) of neural networks. It scans the search space by modifying the The study by Sexton et al. [21] shows that Genetic Algorithm outperforms state of the art techniques like backpropagation in cases where the error surfaces are complex and nonlinear. Other studies (Miller et al. [15], Ding et al. [5]) also show that neural networks have been successfully trained using genetic algorithms as the search strategy.
- Secondly, we try to evaluate if this architecture can achieve the same level of performance as other popular neural network models. For this, we perform experiments on four selected datasets with five different numbers of nodes in the hidden layer. A fully connected neural network with only one hidden layer has been used for the study. It can be extended to architecture of any size. The DBNN model has been compared with: (a) Genetic Algorithm trained neural network, (b) traditional backpropagation model and the models similar to DBNN which were discussed earlier – (c) Bayesian Neural Networks and (d) HyperNetworks.

## 2. Distribution Based Neural Networks (DBNN)

### 2.1. The Principle

Traditional neural networks (TNN) are fully connected networks with each connection represented by weights. They have a complete weight matrix architecture, meaning each connection is associated to its corresponding weight value. While training a neural network model, we train these weights such that the error function at the final layer is optimized. The size of the weight matrix is dependent on the architecture of the network used - number of hidden layers and nodes. If we can decrease the size of the weight matrix by approximating the weights by a distribution, we can reduce the number of trainable parameters and the memory required for the training.

#### Assumption

We assume that for every weight matrix *W*_*ijk*_ connecting the *j*^*th*^ node of layer *i* (*n*_*ij*_) and the *k*^*th*^ node of layer *i* + 1 (*n*_*i*+1,*k*_, ∀*k*), there exists a statistical distribution *D*(*p*_1_, *p*_2_, ..*p*_*h*_) with parameters *p*_1_, *p*_2_, ..*p*_*h*_ that can generate these *W*_*ijk*_ or an equivalent set of weights 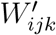, with similar error value on the output layer.

#### Goal of the training

Based on the above assumption, our model determines the parameters *p*_1_, *p*_2_, ..*p*_*h*_ of the distribution instead of the original weight matrix. The *h* values of D are generated by the use of two parameters viz. mean *mu* and standard deviation *sigma* of a Normal distribution, from which individual values *p*_*i*_ are selected by a hard-coded equal probability breakpoint scheme (Section 2.3.1). In other words, we train our neural network to learn the parameters of the distribution.

For a network with *m* features and *n* nodes in the first hidden layer, we can see that the number of parameters for this layer combination is:

TNN: *m* * *n*
DBNN: *m* * 2 (for a distribution with 2 parameters)

Thus, the memory saving is in the order of m *(n-2) for each pair of layers in the network. If we consider a model with 10,000 input features and 10 output classes with 100 nodes in the hidden layer, the savings can be seen in Table 1.

**Table 1:**
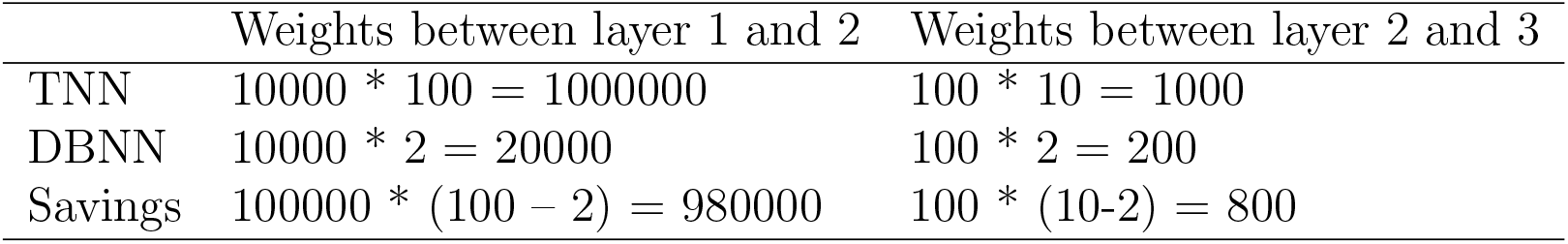
DBNN memory saving for a network with architecture [10000, 100, 10]

We have used Normal distribution for the purpose of our study. However, the model can easily be scaled to new distributions and breakpoint search schemes.

### 2.2. Genetic Algorithms

Genetic Algorithms (GA) is an optimization technique which works on the theory of evolution. The members of population participating in this optimization task are evaluated based on a fitness function. Members with highest fitness are retained in the next generation. New members are created from the previous population by the process of evolution using operations - mutation and crossover. Mutation causes change in some positions of the parent member whereas crossover attaches part of one member to another to form a new member to participate in the next generation. The optimal solution is the one with highest fitness value.

### 2.3. Proposed Architecture of DBNN

Weights of an MLP have been successfully trained by backpropagation and gradient descent for most architectures. Being generated from statistical distribution, the error surface in case of DBNN is presumably highly rugged, poorly differentiable and prone to be trapped in steep local minima. An attempt was made to train DBNN weights using gradient descent optimization with backpropagation of error. The performance observed for this combination of DBNN and GD was not very good compared to other models. As the weight space of the original network is being created through a distribution function and pre-defined breakpoints, in order to find a good solution, the hyperspace of these distribution functions need a wider coverage. To achieve this, Genetic Algorithm was found to be most suitable to train DBNN.

#### 2.3.1. Generative model

A fully connected Neural network can be represented as seen in Fig. 1. Outgoing weights from each node of layer *i* connect the same number of nodes of the *i* + 1^*th*^ layer. We propose to generate equidistant quantiles spaced at a distance of 1/*n* from each other, where *n* is the number of nodes in the *i* + 1^*th*^ layer. The value of z corresponding to *P* (*Z* ≤ *k/n*) ∀ *k* ≤ *n*, can be transformed back to X domain to get the value of each weight

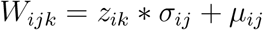

**Figure 1:**
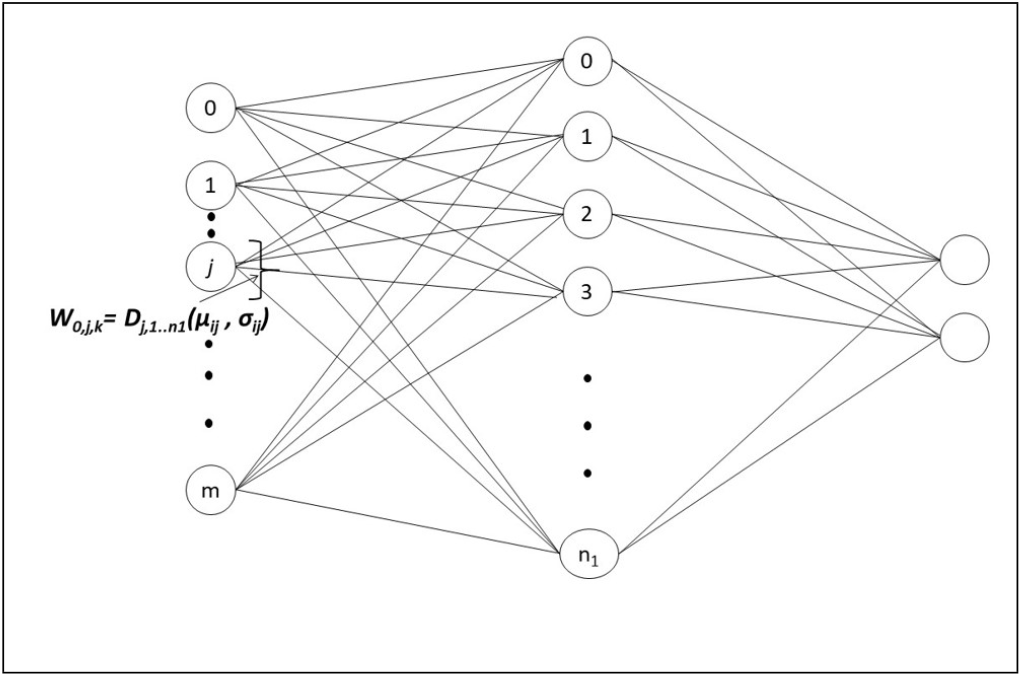
Schematic representation of a prototype DBNN. Every node in the *i*^*th*^ layer is connected to every node in the (*i* + 1)^*th*^ layer through *n*_*i,i*+1_ weights defined by a breakpoint function D on a Normal distribution with trainable mean (*μ*) and standard deviation (*σ*). Weights belonging to first node of the network have been illustrated. Each such matrix has its own distribution hyper-parameters *mu* and *sigma* but all weight generators have an identical breakpoint function, based on quantile values of cumulative frequency distribution.

In the neural network, during forward propagation we take the dot product of input feature values *F*_*i*_ with the matrix *W*_*i*_ to get the features 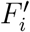 for the next layer. These are then passed through the activation function to get *F*_*i*+1_. For each layer *i*, 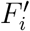 can be calculated as the dot product (•)

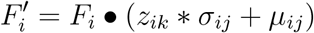

On performing batch operation for each layer *i*,

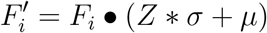

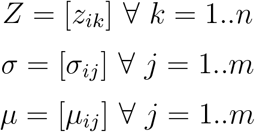

We can see that the dimensions of each matrix are aligned to calculate dot products.

### 2.4. Training the network

In our implementation of DBNN, each generation creates 10 parent networks, out of which the best 5 are used in the next generation, based on a fitness function. The fitness function used here is based on Cross Entropy Loss. Other 5 children are created by operations of crossover and mutation. Single point crossover and random mutation are used in the study, with mutation percent set to 25%. Using these fixed GA parameters, experiments have been conducted with multiple models, with different numbers of hidden nodes. Experiments have also been carried out with four other combinations of the number of parents and parents mating to form the next generation and similar results have been observed for them.

In case of DBNN, while forward propagating the inputs for each neural network, the generative model discussed in Section 2.3.1 is used and then activation values are calculated. The *tanh* function has been used for activation of the hidden layers and *softmax* as the output activation. The weights represented by distribution *D*(*μ*; *σ*) are trained here, as per our assumptions in Section 2.1. Theoretically, it is evident that the number of parameters undergoing optimization to reach the minima is much less in case of DBNN as compared to any traditional neural networks. This makes the model scalable for networks of large size. Without loss of generality, it can be said that the complexity of the algorithms is proportional to the number of generations taken for training and the number of parameters to be trained. It has been shown in the next section that the algorithmic complexity of DBNN is lower than other conventional MLP models trained similarly.

## 3. Experiments Performed

### 3.1. Performance of proposed model

Based on the proposed model of DBNN, the performance has been evaluated on four selected datasets. The datasets were selected to represent both small and moderately large sizes of feature space instance space.

#### Datasets

Four binary classification datasets have been selected from UCI machine learning repository and Kaggle. Categorical labels have been converted to numerical values and all the datasets have been normalized using a min-max scaling. The comparative description of the datasets has been listed in Table 2

1. **Breast Cancer Gene Expression Profiles Dataset** (https://www.kaggle.com/raghadalharbi/breast-cancer-gene-expression-profiles-metabric): This dataset was published by [17] and it contains the rna mutation profiles of breast cancer patients. The target variable is overall survival.
2. **Breast Cancer Wisconsin (Diagnostic) Dataset** (https://archive.ics.uci.edu/ml/datasets/breast+cancer+wisconsin+(diagnostic)): This dataset published by [2] describes the characteristics of the nucleus of a cell extracted from the breast mass. It consists of numerical features corresponding to digitized image of the breast mass. The target variable of is diagnosis of the breast cancer to be malignant or benign.
3. **Parkinsons Dataset** (https://archive.ics.uci.edu/ml/datasets/parkinsons): Parkinsons disease is a neurological disorder which affects movement. The parkinsons dataset was published by [19] and it has features corresponding to voice measurement.
4. **Eye Detection EEG Dataset** (https://www.kaggle.com/c/eyedetection/data): The dataset contains EEG data of 9998 patients captured using EMOTIV EPOC+ technique. It was published by [18] in 2013. MLP techniques have not given good results on this dataset. We have included this to see the performance of DBNN on it.

**Table 2:**
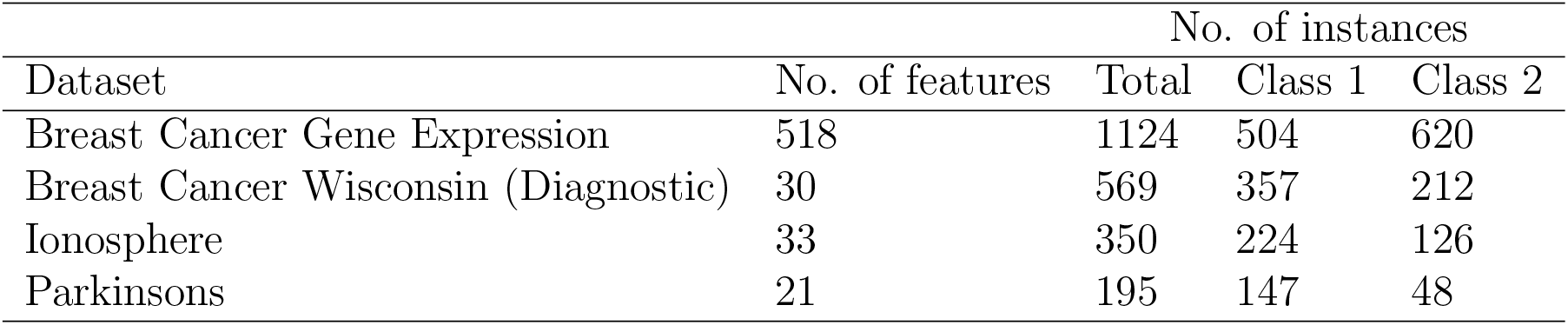
Datasets used in the study

| Dataset | No. of features | No. of instances |  |  |
| --- | --- | --- | --- | --- |
|  |  | Total | Class 1 | Class 2 |
| Breast Cancer Gene Expression | 518 | 1124 | 504 | 620 |
| Breast Cancer Wisconsin (Diagnostic) | 30 | 569 | 357 | 212 |
| Ionosphere | 33 | 350 | 224 | 126 |
| Parkinsons | 21 | 195 | 147 | 48 |

### 3.2. Comparison with other conventional methods

The performance of DBNN has been compared with the following State of the Art models which are similar to DBNN in some way:

1. Genetic Algorithm Neural Networks (GANN)
2. Backpropagation with Gradient Descent
3. Bayesian Neural Networks
4. HyperNetworks

## 4. Implementation

Programming language and packages:

- Pytorch 1.10.1 and Python 3.8.8 have been used in the anaconda3 environment.
- Scikit-learn scientific library has been used to calculate the performance metrics.
- PyGAD, a library for implementing Genetic Algorithms and training artificial neural networks) has been used to implement GANN. DBNN has been implemented by using the PyGAD library.

## 5. Results

### 5.1. Evaluation of DBNN

To perform unbiased evaluation of the models, each dataset has been evaluated for 10 sets of randomly bootstrapped training and test data in the ratio 80:20. Each such dataset has been evaluated for 10 sets of random weights. We know that MLPs can be used to solve any problem by using a single hidden layer architecture, although the number of hidden nodes may become infeasibly large [7]. Our experiment has been conducted with a single hidden layer and extended to different hidden node configurations - [10,20,50,100,200]. The complete scores from the performed experiment can be found in the supplementary information. All reported values are on the test data selected randomly. The average scores have been reported here.

The models have been evaluated under two heads:

1. Performance of the models using evaluation metrics Accuracy and AUC over unseen test data. Table 3 gives the outcome of the experiment for the datasets under study.
2. Complexity of models with respect to memory required to store the parameters and number of generations required to achieve the desired level of change in average error. The last evaluation metric is algorithmic complexity, which is calculated as the number of generations taken for training times the number of parameters trained. Table 4 gives the outcome of the experiment for the four datasets under study with respect to complexity measures.

**Table 3:** Performance Evaluation of Distribution Based Neural Network(DBNN)

| Dataset | Metrics | Nodes in Hidden Layer |  |  |  |  |
| --- | --- | --- | --- | --- | --- | --- |
|  |  | 10 | 20 | 50 | 100 | 200 |
| Breast Cancer | Accuracy | 0.69 | 0.71 | 0.72 | 0.69 | 0.7 |
| Gene Expr | AUC | 0.75 | 0.77 | 0.76 | 0.75 | 0.76 |
| Breast Cancer | Accuracy | 0.94 | 0.95 | 0.95 | 0.96 | 0.95 |
| Wisconsin (D) | AUC | 0.99 | 0.99 | 0.99 | 0.98 | 0.99 |
| Parkinsons | Accuracy | 0.89 | 0.88 | 0.87 | 0.88 | 0.86 |
|  | AUC | 0.83 | 0.85 | 0.85 | 0.84 | 0.84 |
| Eye detection | Accuracy | 0.66 | 0.66 | 0.67 | 0.68 | 0.67 |
|  | AUC | 0.67 | 0.67 | 0.69 | 0.69 | 0.68 |

**Table 4:** Complexity Evaluation of Distribution Based Neural Network(DBNN)

| Dataset | Metrics | Nodes in Hidden Layer |  |  |  |  |
| --- | --- | --- | --- | --- | --- | --- |
|  |  | 10 | 20 | 50 | 100 | 200 |
| Breast Cancer | Generation | 392 | 436.6 | 433.75 | 389.25 | 461.5 |
| Gene Expr | Memory Required | 1014 | 1034 | 1094 | 1194 | 1394 |
| | Algo Complexity ($\times 10^4$) | 39.75 | 45.14 | 47.45 | 46.48 | 64.33 |
| Breast Cancer | Generation | 245.6 | 221.9 | 221.9 | 224 | 216.4 |
| Wisconsin (D) | Memory Required | 80 | 100 | 160 | 260 | 460 |
| | Algo Complexity ($\times 10^4$) | 1.96 | 2.22 | 3.55 | 5.82 | 9.95 |
| Parkinsons | Generation | 221 | 217.7 | 178.1 | 145.7 | 85.1 |
|  | Memory Required | 64 | 84 | 144 | 244 | 444 |
| | Algo Complexity ($\times 10^4$) | 1.41 | 1.83 | 2.56 | 3.56 | 3.78 |
| Eye detection | Generation | 510.5 | 482 | 501.5 | 454.4 | 435.2 |
|  | Memory Required | 48 | 68 | 128 | 228 | 428 |
| | Algo Complexity ($\times 10^4$) | 2.45 | 3.28 | 6.42 | 10.36 | 18.63 |

### 5.2. Comparison of DBNN with other models

Figures 2 and 3 show the comparison of DBNN with other models over Test AUC Score and Algorithmic Complexity. None of the models performs consistently over all types of datasets. To see the overall performance, the average rank of each dataset has been calculated in Table 5. We can see that the average performance of DBNN is second to backpropagation and better than the other three models.

**Figure 2:**
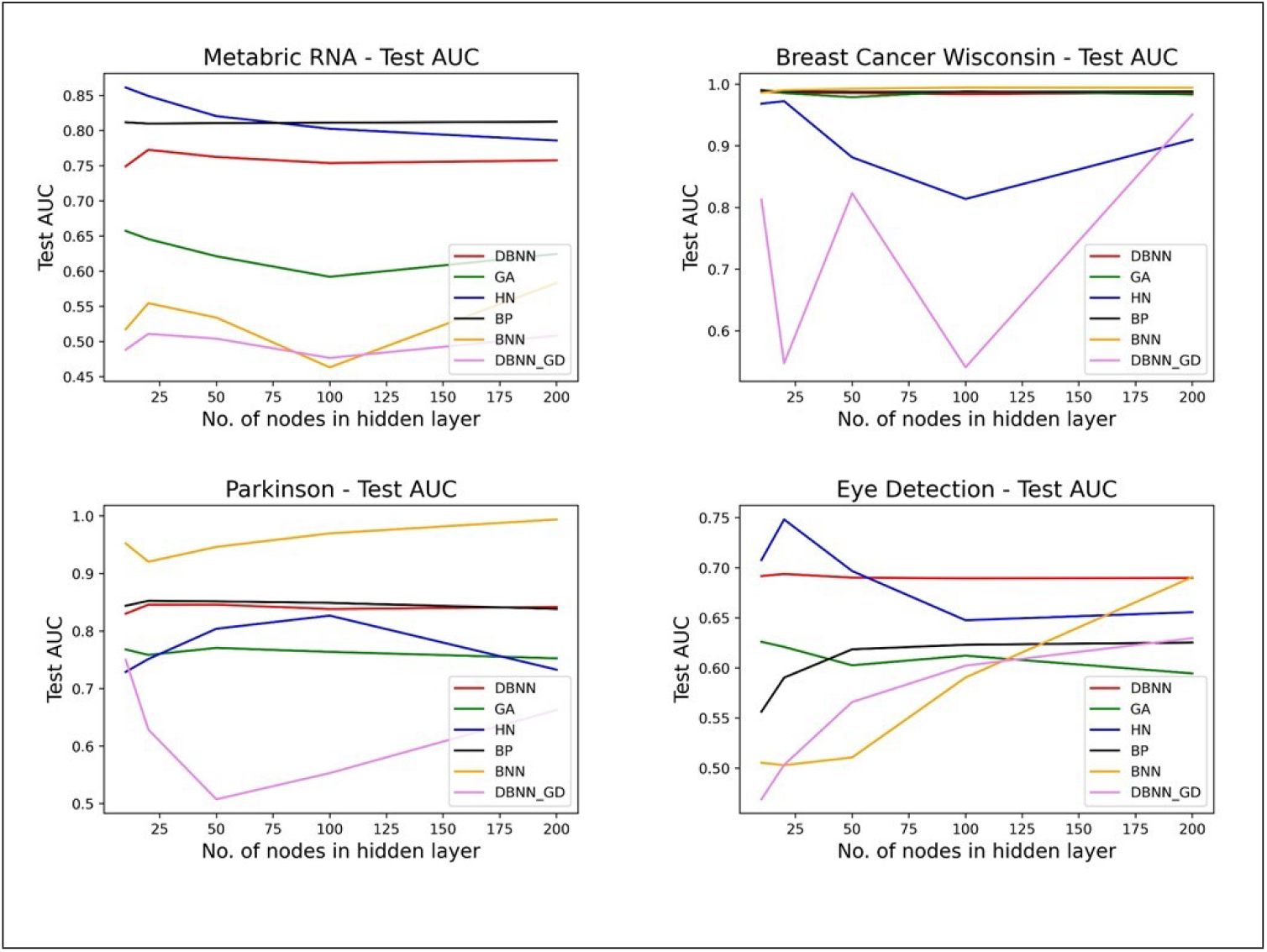
Comparison of the models over Test AUC Score

**Figure 3:**
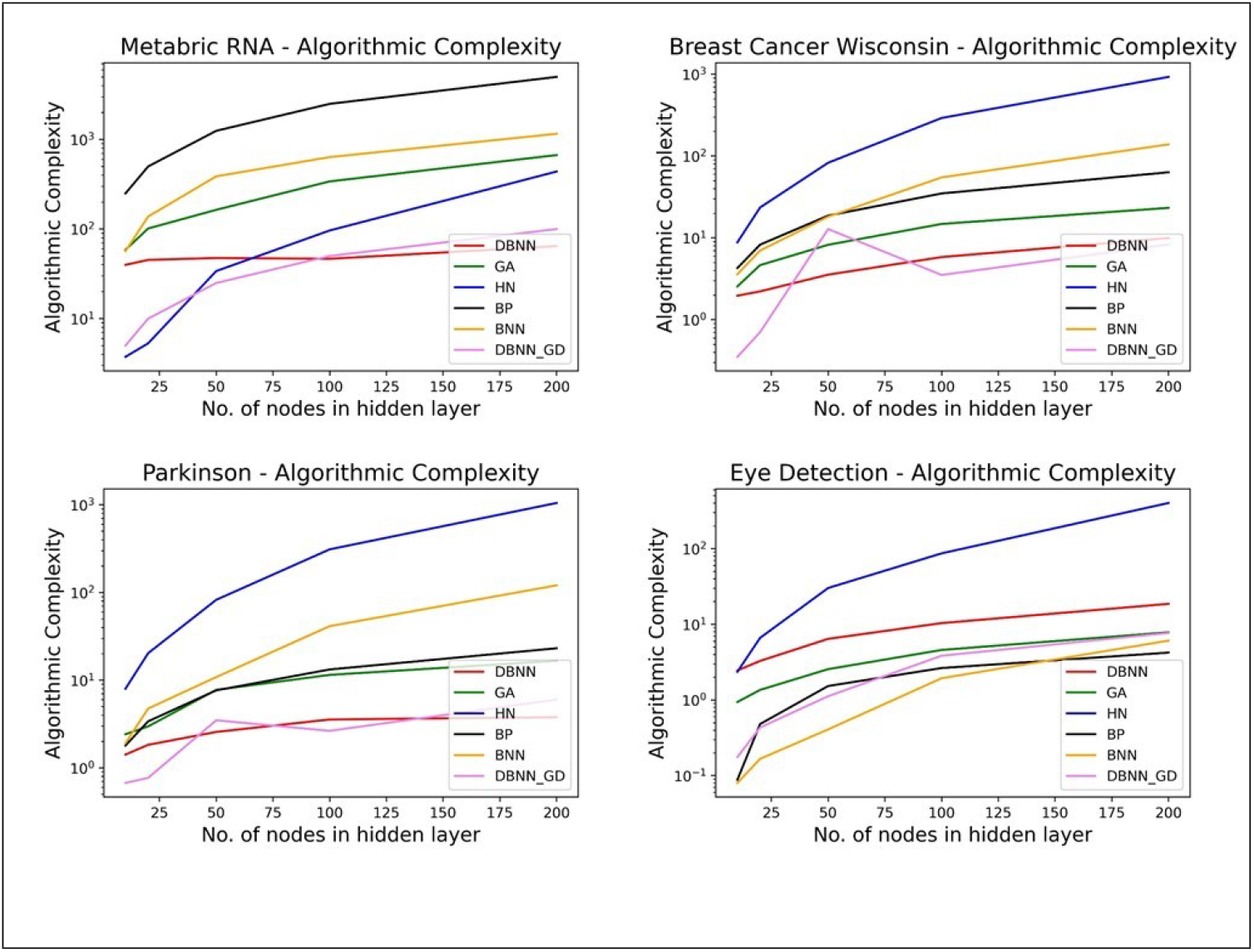
Comparison of the models over Algorithmic Complexity

**Table 5:** Average rank of all models over each dataset with respect to the Test AUC. Lower value of rank corresponds to better performance.

| Dataset/Model | DBNN | BP | BNN | GANN | HN | DBNN-GD |
| --- | --- | --- | --- | --- | --- | --- |
| MetabricRNA | 3 | 1 | 5 | 4 | 2 | 6 |
| Breast Cancer Wisconsin | 3 | 2 | 1 | 4 | 5 | 6 |
| Parkinsons | 3 | 2 | 1 | 5 | 4 | 6 |
| Eye Detection | 1 | 3 | 6 | 4 | 2 | 5 |
| **Average Rank** | 2.5 | 2 | 3.25 | 4.25 | 3.25 | 5.75 |

## 6. Discussion

In this work, we have proposed an alternative method of defining an MLP which uses much less memory than the currently used models. We demonstrated that the proposed model is feasible as it could be trained to achieve comparable performance to the other models. The advantages of the proposed DBNN model are:

- DBNN uses a smaller set of trainable parameters (and less memory) and consistently has least algorithmic complexity while achieving performance comparable to other models.
- HyperNetworks is able to achieve high training accuracy, but for the unseen test set, the accuracy reported is less. Compared to this DBNN is able to achieve better test accuracy, despite lower training accuracy than HyperNetworks, suggesting that it may be able to avoid overfitting better.
- For the EEG eye detection dataset, the authors note in their paper that traditional MLP training algorithms do not perform well on the data. In our experiment we find that DBNN performs better than other models by achieving a maximum test accuracy of 67.57% with AUC of 0.69 as compared to the average accuracy of 64.87% and average AUC of 0.62 across other models. This amounts to a substantial advantage of 7 and 2.5 percentage points in AUC and in accuracy respectively.
- As DBNN is not trained using backpropagation, it is not limited to differentiable activation functions such as sigmoid or arctan. Although, we have not experimented extensively on this aspect in the current work, we believe, use of novel activation functions coupled with DBNN model, will render the model free from currently recognized problems of MLP training such as vanishing gradient, oscillating error function and local non-optimal minima.
- Furthermore, since the actual training parameters are fewer, the model is expected to have better generalisation and less likely to get overfitted, as mentioned on specific datasets above. Whether this comes with a loss of trainability in some situations is yet to be investigated and will be revealed over the subsequent works on the subject. At this point, all the four data sets could be trained very well at high levels of accuracy, which gives confidence in its huge potential.
- Also, it is obvious that DBNN complexity gain is most visible when very long hidden layers are to be used. Thus, another issue to be investigated in the future is whether a shallow and very wide MLP model can reach better performances on this model, which can potentially outperform deeper models.

Thus, the current work provides a framework to explore many other possibilities, while at the outset establishing its performance and viability in selected examples.

## 7. Conclusion

A neural network model, DBNN in which statistical distributions are used to initialise and train connection weights is proposed. Evaluation of the DBNN on selected datasets prove that such models can not only achieve similar performance as the fully defined MLPs with a much reduced memory complexity, but may also outperform them in terms of generalisation due to lower number of parameters to be trained. DBNN could be trained using Genetic algorithm effectively in each of the 4 examples tested, suggesting its broader applicability. We believe that DBNN based models will provide a powerful alternative architectural, initialization and training frameworks for neural networks and hold the promise of outperforming currently known MLP models.

## Notes

### Competing Interest Statement

The authors have declared no competing interest.

## References

[1] Amato, F., López, A., Peña-Méndez, E.M., Vaňhara, P., Hampl, A., Havel, J., 2013. Artificial neural networks in medical diagnosis.

[2] Bennett, K.P., Mangasarian, O.L., 1992. Robust linear programming discrimination of two linearly inseparable sets. Optimization methods and software 1, 23–34.

[3] Besari, A.R.A., Prabuwono, A.S., Zamri, R., Palil, M.D.M., 2010. Computer vision approach for robotic polishing application using artificial neural networks, in: 2010 IEEE Student Conference on Research and Development (SCOReD), IEEE. pp. 281–286.

[4] Courbariaux, M., Bengio, Y., David, J.P., 2015. Binaryconnect: Training deep neural networks with binary weights during propagations. arXiv preprint arXiv:1511.00363.

[5] Ding, S., Su, C., Yu, J., 2011. An optimizing bp neural network algorithm based on genetic algorithm. Artificial intelligence review 36, 153–162.

[6] Glorot, X., Bengio, Y., 2010. Understanding the difficulty of training deep feedforward neural networks, in: Proceedings of the thirteenth international conference on artificial intelligence and statistics, JMLR Workshop and Conference Proceedings. pp. 249–256.

[7] Goodfellow, I., Bengio, Y., Courville, A., Bengio, Y., 2016. Deep learning. volume 1. MIT press Cambridge.

[8] Ha, D., Dai, A., Le, Q., 2016. Hypernetworks.

[9] Jesus, R., Antunes, M., da Costa, R., Dorogovtsev, S., Mendes, J., Aguiar, R., 2020. Effect of the initial configuration of weights on the training and function of artificial neural networks. arXiv preprint arXiv:2012.02550.

[10] Kim, Y., Yun, W.J., Lee, Y.K., Jung, S., Kim, J., 2021. Trends in neural architecture search: Towards the acceleration of search, in: 2021 International Conference on Information and Communication Technology Convergence (ICTC), IEEE. pp. 421–424.

[11] Kingma, D.P., Salimans, T., Welling, M., 2015. Variational dropout and the local reparameterization trick. arXiv preprint arXiv:1506.02557.

[12] MacKay, D.J., 1992. A practical bayesian framework for backpropagation networks. Neural computation 4, 448–472.

[13] Martens, J., 2010. Deep learning via hessian-free optimization., in: ICML, pp. 735–742.

[14] Mas, J.F., Flores, J.J., 2008. The application of artificial neural networks to the analysis of remotely sensed data. International Journal of Remote Sensing 29, 617–663.

[15] Miller, G.F., Todd, P.M., Hegde, S.U., 1989. Designing neural networks using genetic algorithms., in: ICGA, pp. 379–384.

[16] Nanda, S.K., Panda, S., Subudhi, P.R.S., Das, R.K., 2012. A novel application of artificial neural network for the solution of inverse kinematics controls of robotic manipulators. International Journal of Intelligent Systems and Applications 4, 81–91.

[17] Pereira, B., Chin, S.F., Rueda, O.M., Vollan, H.K.M., Provenzano, E., Bardwell, H.A., Pugh, M., Jones, L., Russell, R., Sammut, S.J., et al., 2016. The somatic mutation profiles of 2,433 breast cancers refine their genomic and transcriptomic landscapes. Nature communications 7, 1–16.

[18] Rösler, O., Suendermann, D., 2013. A first step towards eye state prediction using eeg. Proc. of the AIHLS 1, 1–4.

[19] Sakar, C.O., Serbes, G., Gunduz, A., Tunc, H.C., Nizam, H., Sakar, B.E., Tutuncu, M., Aydin, T., Isenkul, M.E., Apaydin, H., 2019. A comparative analysis of speech signal processing algorithms for parkinson’s disease classification and the use of the tunable q-factor wavelet transform. Applied Soft Computing 74, 255–263.

[20] Saxe, A.M., McClelland, J.L., Ganguli, S., 2013. Exact solutions to the nonlinear dynamics of learning in deep linear neural networks. arXiv preprint arXiv:1312.6120.

[21] Sexton, R.S., Dorsey, R.E., Johnson, J.D., 1998. Toward global optimization of neural networks: A comparison of the genetic algorithm and backpropagation. Decision Support Systems 22, 171–185.

[22] Shayer, O., Levi, D., Fetaya, E., 2017. Learning discrete weights using the local reparameterization trick. arXiv preprint arXiv:1710.07739.

[23] Tkáč, M., Verner, R., 2016. Artificial neural networks in business: Two decades of research. Applied Soft Computing 38, 788–804.

